# CellsFromSpace: A fast, accurate and reference-free tool to deconvolve and annotate spatially distributed Omics data

**DOI:** 10.1101/2023.08.30.555558

**Authors:** Corentin Thuilliez, Gael Moquin-Beaudry, Pierre Khneisser, Maria Eugenia Marques Da Costa, Slim Karkar, Hanane Boudhouche, Damien Drubay, Baptiste Audinot, Birgit Geoerger, Jean-Yves Scoazec, Nathalie Gaspar, Antonin Marchais

## Abstract

Spatial transcriptomics involves capturing the transcriptomic profiles of millions of cells within their spatial contexts, enabling the analysis of cell crosstalk in healthy and diseased organs. However, spatial transcriptomics also raises new computational challenges for analyzing multidimensional data associated with spatial coordinates.

In this context, we introduce a novel framework called CellsFromSpace. This framework allows users to analyze various commercially available technologies without relying on a single-cell reference dataset. Based on the independent component analysis, CellsFromSpace decomposes spatial transcriptomic data into components that represent distinct cell types or activities. Here, we demonstrate that CellsFromSpace outperforms previous reference-free deconvolution tool in term of accuracy and speed, and successfully identify spatially distributed cells as well as rare diffuse cells on datasets from the Visium, Slide-seq, MERSCOPE, and COSMX technologies.

The framework provides a user-friendly graphical interface that enables non-bioinformaticians to perform a full analysis and to annotate the components based on marker genes and spatial distributions. Additionally, CellsFromSpace offers the capability to reduce noise or artifacts by component selection and supports analyses on multiple datasets simultaneously.

**Graphical abstract:** 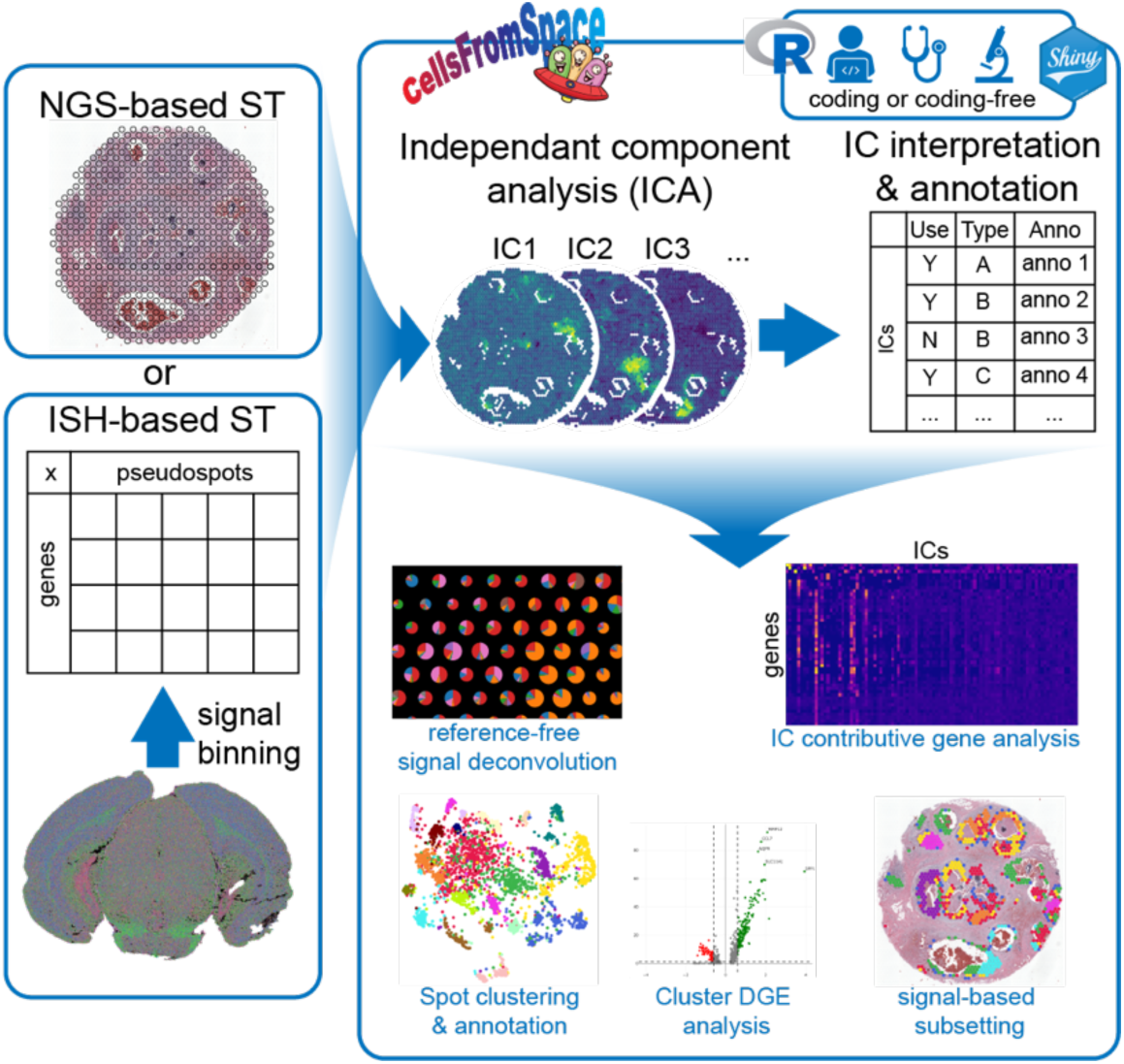

## Introduction

Spatial transcriptomics (ST) have emerged as some of the most promising technologies to analyze spatial distribution and context of cell types and activities within tissues^1^. Commercially available technologies are divided into two main categories: spatially barcoded next generation sequencing-based (10X Visium^1^, Slide-seqV2 now Curio Seeker^2^, Stereo-seq^3^) and transcript level panel-based high throughput in situ hybridization (ISH) or sequencing (ISS) approaches (Vizgen MERSCOPE^4^, Nanostring CosMX^5^, 10X Xenium^6^). The former enables whole or panel transcriptome analysis on tissue sections at varying degrees of resolution, from subcellular to quasi cellular resolution, depending on the technology. These methods generate highly dimensional, sparse, spatially distributed data, which can be demanding on computational resources, and still requires the development of new algorithms to exploit jointly the molecular data, spatial coordinates and tissue images^7^.

10X Genomic’s Visium technology, currently the most widespread commercial solution by publication metrics, outputs manageable data sizes at the cost of lower resolution compared to other techniques, with 55µm diameter spots typically encompassing 1-20 cells. Due to the “mini-bulk” nature of the technology, classical methods used in single cell analysis are less efficient and will cluster spots according to the cell mixture. In the worst-case scenario, such as the complex and unevenly spatially distributed tumor tissues, numerous clusters will be generated obfuscating the cell composition of each spot. Therefore, a deconvolution/decomposition step is often required to gain insight into the mixture of cells populating each spot. These deconvolution methods, at the exception of few^8^, are mostly scRNA-seq reference based^9,10–19^, with known drawbacks such as the necessity for high-quality reference datasets and the loss of information.

Recently, Miller et al^8^, with STdeconvolve, proposed a reference free method to decompose the signal with Latent Dirichlet Allocation (LDA), a text processing technique, approximating genes to words and spot to documents. Their work presents several interesting developments but the known drawbacks of LDA (Sensitivity to noisy and sparse data – Difficulty to detect rare topics – Fixed numbed of topics) might limits its performance when applied to ST dataset. Here, we propose a new reference free decomposition framework for ST dataset, named CellsFromSpace (CFS) that overcomes some of the limitation of LDA. CFS is based on the independent component analysis (ICA), a blind source separation technique that attempts to extract signal sources from a mixture of these sources^20^. ICA has been successfully applied to bulk transcriptomic data in hundreds of publications and shows the best performance over other methods to detect gene module from bulk RNA-seq^21^. ICA performance is directly dependent on the ratio between the number of sources and number of sampled mixtures. With thousands of spots (mixtures) each covering few cell types (Sources), we assumed that ICA should perform well to decompose cell types and activities from ST data. Additionally, through the biological interpretation and spatial distribution analysis of the independent components (ICs), an expert in the field can supervise the removal of noise and artifacts, as demonstrated in electroencephalography (EEG), functional magnetic resonance imaging (fMRI), and recent advancements in neuroscience^22,23^. Hence, in ICA latent space of ST data one can select, isolate, or remove background noise and parasitic signals, such as cell death, that are not relevant for downstream analysis, without altering the signal of interest. Finally, the permissiveness of ICA regarding over-decomposition^24^ is a significant advantage that allows to fix an arbitrary large number k with a minimum drawback of generating near identical components, easily corrected by component annotations and subsequent merging. Altogether, the characteristics of ICA in combination to an expert-supervised annotation and component selection, implemented in the CFS framework, enables: i) to avoid dimensionality optimization steps, ii) to select biologically relevant components and to remove noise, iii) to study specific cell subtypes in the IC latent space, and iv) to identify conserved signal though multi-sample ST datasets.

When applied to ST analysis, CFS allows a biologically relevant decomposition of the spot mixture into its subcomponents responsible for common expression patterns observed within the tissue, such as cell type signatures, biological processes and tissue organization.

CFS consists of a preprocessing pipeline and an easy-to-use Shiny user interface (UI) to analyze, annotate, visualize, and subset ST data. CFS was used to fully process sample datasets from multiple major sequencing-based ST technologies and was also applied to ISH ST as an intermediary tool for sample screening and identification of regions of interest. This highlights the flexibility and effectiveness of a semi-supervised ICA-based signal decomposition for the analysis of ST in healthy and diseased samples. The companion shiny interface enables non-bioinformaticians to quickly perform, without programming, a complete analysis of ST data from Visium, Slide-seq, MERSCOPE or CosMX dataset; and to generate Seurat^25^ objects compatible with subsequent analyses.

## Methods

### CFS overview and use case

The package is split into two main components: A) the package itself which includes functions to preprocess data loaded in *Seurat* by i) *prepare_data* which normalizes the count matrix, ii) *RunICA* which runs the *fastica* algorithm to separate the signal into independent components (IC), corrects the signs of ICs (by convention we consistently flip the sign to define the max weight as positive), and filters out ICs with a kurtosis below a user-defined threshold (default: 3), iii) *Enrich_ICA* which queries EnrichR databases to run and curate functional enrichment analyzes for individual ICs, and iv) in addition, functions to convert ISH ST data into a format compatible with CFS, and B) a Shiny UI to run the previously mentioned preprocessing steps and downstream annotation and analysis of ICA results. A critical step of CFS’s analytical workflow is the manual annotation of ICs by scientists. To improve the ease, speed, and efficacy of this critical step by biologists, clinicians, or any experts, CFS’s Shiny UI provides a series of panels to visualize different aspects of the ICs such as: i) global gene x IC heatmap of the top contributing genes for all ICs, ii) spatial distribution of ICs and their contributing genes, iii) IC-specific gene x IC and gene x spot heatmaps showing the contribution of IC-defining genes in ICs and cells, and iv) annotated bar graph visualization of functional enrichment analyses from EnrichR of the contributing genes for each ICs.

Once annotated, relevant ICs can be used to calculate spot clustering and UMAP dimensionality reduction. Marker genes for the calculated clusters can also be calculated within the Shiny UI. However, an advantage of ICA, that directly captures cell type signatures, is the ability to remain in the variable latent space. To interpret the mixture in IC space of each spot, CFS provides spatial and UMAP scatter pie chart representations of data. These scatter pie representations allow the observation of all or a selected subset of ICs weight on each spot as well as an annotation-based categorical representation of these ICs. Due to the assumptions of ICA, scaling between ICs is not meaningful. Therefore, it is important to note that scatterpie does not represent proportion itself. Instead, for each IC, it represents the centered weight among all the spots. We believe that this limitation does not hinder the CFS analysis. In our opinion, when dealing with spots of 55μm spaced 110μm apart (and thus lacking information about neighboring cells), identifying the presence of cells is more relevant than inferring their exact composition.

The Shiny UI allows for a complete analysis workflow from the loading of SpaceRanger output to the easy exporting of publication-ready figures of all visualizations in png, jpeg, pdf or svg format. Data generated within the application can also be exported in rds format integrally or subsetted from within the tool. The resulting *Seurat* objects containing all curated annotations, calculated metadata and dimensionality reductions can be loaded directly in R for more complex downstream analyses with compatible workflows.

### Samples used

The package efficiency was assessed using Visium reference sample datasets of Mouse Brain Coronal Section, Human Breast Cancer and Human Prostate Cancer. Slide-seqV2 mouse hippocampus sample dataset was obtained through the *SeuratData* package. MERSCOPE sample dataset MERFISH Mouse Brain Receptor Map was obtained from Vizgen (https://info.vizgen.com/mouse-brain-data). CosMX sample datasets and annotations of formalin-fixed paraffin-embedded (FFPE) human non-small cell lung cancer were obtained from nanostring (https://nanostring.com/products/cosmx-spatial-molecular-imager/ffpe-dataset/)

### Preprocessing pipeline

All samples were pre-processed using the CFS package and the following pipeline. Samples were normalized using *Seurat*’s *sctransform* function. Using the *ICASpatial* function, 100 ICs were calculated for the ICA analysis with 600 iterations using the *Icafast* method. Only leptokurtic ICs (kurtosis > 3) were retained for downstream processing and analysis. By convention, IC sign correction was then applied by flipping signs to define max IC weights as positive. The sign doesn’t alter the interpretation of ICs, but we have empirically observed that this correction improves the interpretation of ICs by experts and better fit with the biology. Functional enrichment analysis of IC-contributing genes (defined as genes with feature loading absolute z-score ≥ 3) was done using the *enrichR* package’s *Show_Enrich* function for the desired databases

### Pseudotime analysis

Pseudotimes, branches and diffusion maps were computed from the independent components annotated as tumor, only for the spot overlapping tumor cells, using the corresponding functions of the destiny 2.0 package^26^.

### ISH data pseudospot binning

To process ISH technologies’ datasets within CFS, samples first needed to be converted into a pseudo-spot format. To do so, a count matrix was recreated using the reported detected transcript table from standard output using CFS’s *Create_vizgen_seurat* or *Create_CosMX_seurat* functions (detected_transcript.csv for MERSCOPE and tx_file.csv for CosMX) with transcripts placed in grids of variable bin sizes (40×40 µm for MERSCOPE, 200×200 px or 24×24 µm for CosMX). Each pseudo-spot of this grid was then treated similarly to a Visium spot with the corresponding associated transcript pseudo-counts. A *Seurat* object was then created using this matrix as count matrix input and the pre-processing pipeline was run as described previously. For the MERSCOPE sample, pseudospots with less than five total transcripts detected were filtered out.

### Comparative performance with STDeconvolve

Visium Mouse brain and breast cancer and CosMX NSCLC datasets were deconvoluted using the standard STdeconvolve workflow. Briefly, all pseudospots previously filtered by feature count were processed. Genes used for the analysis were restricted using *restrictCorpus* function to restrict over-dispersed genes. Genes above 5% and under 100% of pseudospots were thus included. Latent Dirichlet allocation (LDA) was applied to find K latent topics. For each sample, the K value with minimum perplexity was kept: 38 for 10x mouse brain, 30 for 10x breast cancer, 22 for 10x prostate cancer, and 57 for integrated CosMX NSCLC.

Comparative annotation performance assessment was conducted first by extracting ground truth cell annotations from the Giotto-processed object of the NSCLC dataset and attributed by pseudospot. Spatial composition and gene signature correlations were computed using Pearson’s r coefficient. For error calculations, ground truth cell types were mapped to ICs and topics by their gene signature, simulating perfect feature annotation. For this, mean gene expression for each ground truth cell type was correlated using Pearson’s r coefficient with gene weights by feature and z-score was calculated for all features per cell type. Features with z-score values > 2 were annotated and collapsed to the designated cell type. Relative cell type composition for each modality was calculated by pseudospot, with features contributing to <5% being filtered out for Stdeconvolve and CFS. Isomeric log ratio transformation (ILR)^27^ was applied to compositional data and root-mean-square error (RMSE) values calculated with

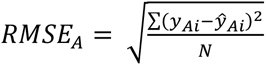

Where *N* is the number of pseudospots, y*Ai* the predicted proportion for ILR dimension A and ŷ*Ai* the ground truth proportion for this dimension. One-tailed Diebold-Mariano test^28^ was used to compare overall RMSE between algorithms.

## Results

### Visium reference data analysis

The efficiency of the CFS workflow was first assessed by analyzing Spatial Gene Expression Visium datasets available from 10x Genomics. Visium fresh frozen and FFPE samples of adult mouse brain and human tumors were thus analyzed using our standard coding-free methodology directly in the Shiny application.

For the FFPE mouse brain sample, after standard preprocessing with CFS (see Methods), 92 ICs passed the kurtosis threshold for downstream analysis. Taking advantage of the annotation tools included in the Shiny app, in half a day, 75 ICs were manually annotated by a biologist as relevant based on their distribution pattern or gene signature (Table S1). Most ICs were directly associated with well-defined brain structures, demonstrating the direct capture by ICs of spatially distributed gene co-expression. Interestingly, ICA also captured ICs associated with diffuse or infiltrating cell populations such as microglia or oligodendrocytes (Fig. 1a) as defined by the contributing genes (Table S2). This confirms the capacity of ICA to isolate cell type specific signal to enable reference-free signal decomposition. Then, from the Shiny app, spot clustering using a Louvain algorithm (resolution = 3.8) based on these 75 dimensions generated 37 clusters closely recapitulating the layers and substructures of the mouse brain as compared to the Allen mouse brain atlas reference for the same coronal layer (position 269, Fig. 1b). For instance, cortical layers or Ammon’s horn pyramidal layer sections are clearly defined, within the limits of Visium’s spot resolution, as distinct clusters since most layers are explained by specific ICs. The distribution of clustered spot in the UMAP embedding (Fig. 1c) demonstrated the unambiguous spot cluster assignment emphasizing the benefits of the expert annotation and the exclusion of ICs considered as noise or specific to one spot.

**Figure 1.**
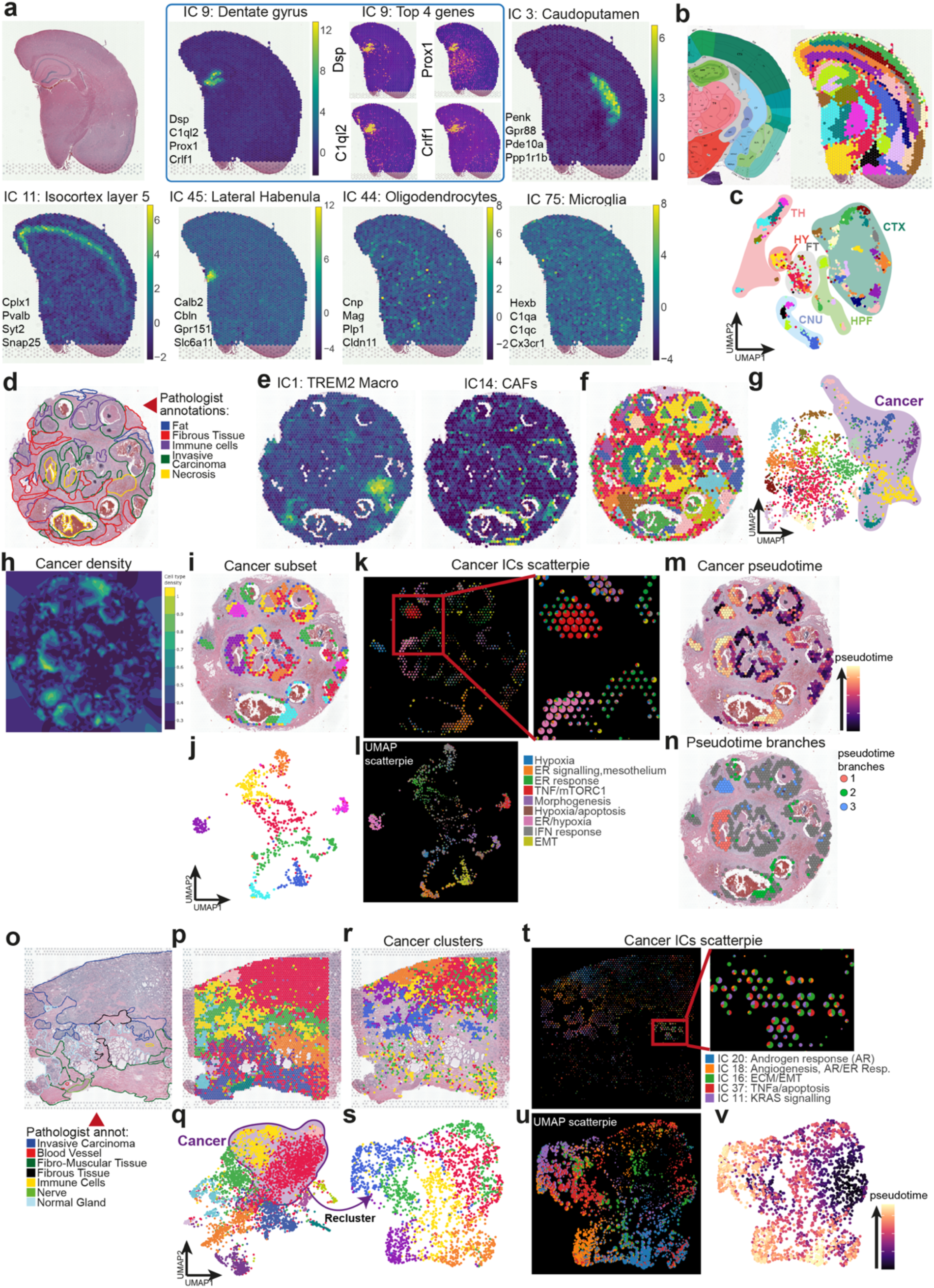
CFS can deconvolute Visium spatial transcriptomics data using independant component analysis. (a-c) Visium FFPE mouse brain analysis using CFS showing (a) H&E reference slide of the sample (top left) and examples of ICs associated to various brain substructures and diffuse cell types each with their top 4 contributive genes (See Supp. Table 2 for full list of contributive genes), and spatial feature example of contributive genes for IC 9: Dentate gyrus. (b) reference slide from the Allen mouse brain Atlas (left) and identified substructures following spot clustering using Louvain algorithm at a 3.8 resolution with filtered and annotated ICs as input (right) showing excellent substructure resolution. (c) 2 dimensional UMAP projection of spots after clustering with corresponding brain zones annotated. CNU: Cerebral nuclei, CTX: Cortex, DG: Dentate gyrus, FT: Fiber tracts, HPF: Hippocampus, HY: Hypothalamus, TH: Thalamus. (d-n) Visium FFPE human breast cancer analysis using CFS. (d) Reference H&E slide with pathologist annotations from 10X Genomics. (e) Sample spatial projection of ICs related to distinct tumor stromal cells (See Supp. Table 4 for full list of contributive genes). Spatial (f) and UMAP (g) projections of spot clustering using Louvain algorithm at a 1.2 resolution with all filtered and annotated ICs as input with cancer-associated clusters highlighted in the UMAP projection. (h) Kernel density projection on the spatial embedding of Cancer-associated signal (based on sum of cancer IC weights, see fig. S1 for tumor stroma components kernels) allowing for automatic subsetting of cancer-associated spots within CFS. Spatial (i) and UMAP (j) projections of cancer spots following manual subsetting within CFS colored after reclustering using Louvain algorithm at a 1.0 resolution with only cancer-associated ICs as input. Scatterpie representation of cancer IC weights in spatial (k) and UMAP (l) projections allows for the rapid visualization of the ICs associated with distinct spot clusters and their respective annotations within CFS. Spatial projection of pseudotime calculation (m) and distinct branches (n) following trajectory inference with Destiny’s DPT algorithm (see fig. S2) showing three clear cancer subpopulations within the sample. (o-v) Visium FFPE human prostate cancer analysis using CFS. (o) Reference H&E slide with pathologist annotations from 10X Genomics. Spatial (p) and UMAP (q) projections of spot clustering using Louvain algorithm at a 0.5 resolution with all filtered and annotated ICs as input with cancer-associated clusters highlighted in the UMAP projection. Spatial (r) and UMAP (s) projections of cancer spots following manual subsetting within CFS colored after reclustering using Louvain algorithm at a 0.5 resolution with only cancer-associated ICs as input. Scatterpie representation of cancer IC weights in spatial (t) and UMAP (u) projections allows for the rapid visualization of the ICs associated with distinct spot clusters and their respective annotations within CFS. UMAP projection of pseudotime calculation (v) following trajectory inference with Destiny’s DPT algorithm (see fig. S3) showing cancer subpopulations within the sample.

Efficient for healthy brain, a well-regionalized organ, CFS capturing both regionalized and isolated cells should fit to medical research projects, where we frequently strive to characterize small groups of cells exhibiting distinct behaviors. This is of particular interest for the analysis of spatially heterogeneous tissues with invasive cells such as tumors. We thus used CFS’ Shiny UI to analyze FFPE breast cancer (Fig. 1d, Table S3-4) and prostate cancer (Fig. 1o, Table S5-6) samples from 10X Genomics to determine the cell type composition and biological activities within spots of tumor tissues. After IC annotation and filtering, 46 ICs (Fig. 1e, Table S3) were used for spot clustering of breast cancer (Figs 1f,g), and 36 ICs for clustering of prostate cancer (figs 1p,q), yielding clusters closely mapping to the pathologist annotations for each tissue. Due to the size of Visium spots, most are composed of multiple cell types in such tissues. 2D embedding and clustering thus tend to be driven by cellular mixture rather than real identity. However, using the interpretable ICA latent space, distinct cell types can directly be mapped both spatially and onto the UMAP embedding. CFS also allows for kernel density mapping of IC categories to easily visualize the distribution of signal associated to broad cell types of interest such as cancer cells (Fig 1h), lymphoid, myeloid or stromal cells within tumors (supp. Fig. 1). This constitutes a semi-supervised strategy implemented within CFS to subset spots of interest with specific annotation. Alternatively, spot subsetting can be done manually based on cluster identity following clustering, for instance to isolate spots with a cancer signature based on IC enrichment (Figs 1i,r). Spot reclustering of the subsetted object using only cancer-associated ICs allows for a finer dissection of cancer phenotypes within samples (Figs 1j,s). To analyze further, CFS integrates multiple visualization tools to interrogate the cellular composition and distribution within samples. For instance, scatter pie representation allows for visual breakdown of cellular composition for particular cell populations based on IC annotation both in spatial (Figs 1k,t) and UMAP (Figs 1l,u) embeddings. The isolation of spots and ICs specific to cell types and their export in a Seurat object (directly from the Shiny app) enables downstream analysis using any other compatible packages. For instance, a trajectory inference analysis of the breast cancerous spots was done using the Destiny package directly on the IC latent space (fig 1m & s2a) and revealed 3 distinct phenotypic branches (Figs 1n, s2b,c). Genes associated with each branch were extracted using *glmnet* (alpha 0.02, Figs s2d,e) and revealed a subpopulation characterized by high IGFBP5, GSTP1 and GNAT3 and low SERF2 expression, a second with high TGM2, SERPINA3, IL32, UBD and ICAM1 and low SCGB1D2, SCGB2A2, AZGP1, MUCL1 and DBI expression, and a third with high SOD2 and low SCGB1D2 AZGP1 and DBI expression.

For the prostate cancer sample, the same trajectory inference methodology (Figs 1u, s3a) also identified 3 branches (Figs s3b,c,d), although less defined, and a *glmnet* analysis revealed the genes associated with these phenotypic paths and distinguished by their expression of ODC1, SPON2, ADGRF1, TSPAN8, CLDN3 and KRT8 among others (Figs s3e,f).

With ICA, CFS efficiently identified distinct cancerous cell phenotypes within a tumor. Furthermore, it identified and distinguished stromal signatures such as immune infiltrates without the use of prior knowledge about the sample or reliance on external reference single cell datasets, which is particularly relevant for highly heterogeneous tissues such as cancer where reference atlases might not faithfully recapitulate the patient’s cancer phenotype or tumor composition.

### Slide-seqV2 data analysis

To demonstrate the compatibility of CFS with other technologies, we analyzed the *ssHippo* mouse hippocampus Slide-seqV2 reference data from the *SeuratData* package using CFS’s standard pipeline. Despite Slide-seqV2 enabling near-single cell resolution (10µm diameter spots), the regular grid of spot still captures mixture of cells. Therefore, ICA should remain very efficient to identify structure and cell type-specific signal. Of the 100 computed ICs, none were thresholded out by kurtosis value and 70 were kept after manual annotation (Table S7-8). Thirty ICs were removed after thorough annotation by a biologist because the signal had no biological meaning (considered as “noise”), or was explained by only one or few spots. The remaining ICs were found to be associated to both A) brain ontologies (ex. IC 8: dentate gyrus granule cell layer, IC 10: Ammon’s horn field 1 pyramidal layer, IC 18: Hippocampal stratum oriens & radiatum and molecular and polymorph layers of the dentate gyrus, IC 37: Ammon’s horn field 2 pyramidal layer and Fasciola cinerea, Fig 2a), and B) distinct cell types (ex. IC 5: ependymal cells, IC 7: ventricular and leptomeningeal cells (VLMC), 7 varieties of neurons, including IC 13: interneurons, and IC 69: proliferating neural stem cells, Fig 2a). Spot clustering based on these ICs allowed for a detailed mapping of the mouse hippocampal region with comparable resolution to the Allen mouse brain atlas reference (Fig. 2b). Indeed, of the 48 recovered clusters obtained with Louvain resolution of 0.95, 45 could be directly annotated based on IC representation (Fig. 2c). Of note, using this approach we were able to successfully identify a cluster of spots (Fig 2c; cluster 14) associated to the small CA2 pyramidal layer (pl) which is typically missed by other algorithms. Once identified, differential expression of CA2pl spots in comparison with CA1pl and CA3pl (clusters 10 and 13 respectively) revealed lower expression levels of calcium channel-related genes such as calmodulin 2 (*Calm2*), ATPase plasma membrane Ca2+ transporting 1 (*Atp2b1*), Protein phosphatase 3 catalytic subunit alpha (*Ppp3ca*), neurogranin (*Nrgn*), protein kinase C beta (*Prkcb*) and an increase in Purkinje cell protein 4 (*Pcp4*) a modulator of calmodulin activity. Interestingly, distinct clusters were found to be associated to either brain ontologies such as the dentate gyrus, 3^rd^ ventricle or thalamus, or to more diffuse cell populations like neurons, microglia, oligodendrocytes or astrocytes. Multiple subclusters for these diffuse populations are often identified with specific transcriptional signatures. Probably, for this diffuse cell population, each spot not covering only a single cell, CFS captured a mixture containing transcripts associated to both a cell and its immediate microenvironment. These results again highlighted the limits of clustering-based methods in sequencing-based ST. Limits circumvented by remaining in the ICA latent space where the IC weight is a direct proxy of the presence of a cell type in each spot. Moreover, these results underlined the complexity of biological tissues, even those as organized as the brain where infiltrating cells are abundant and potentially highly relevant to biologists.

**Figure 2.**
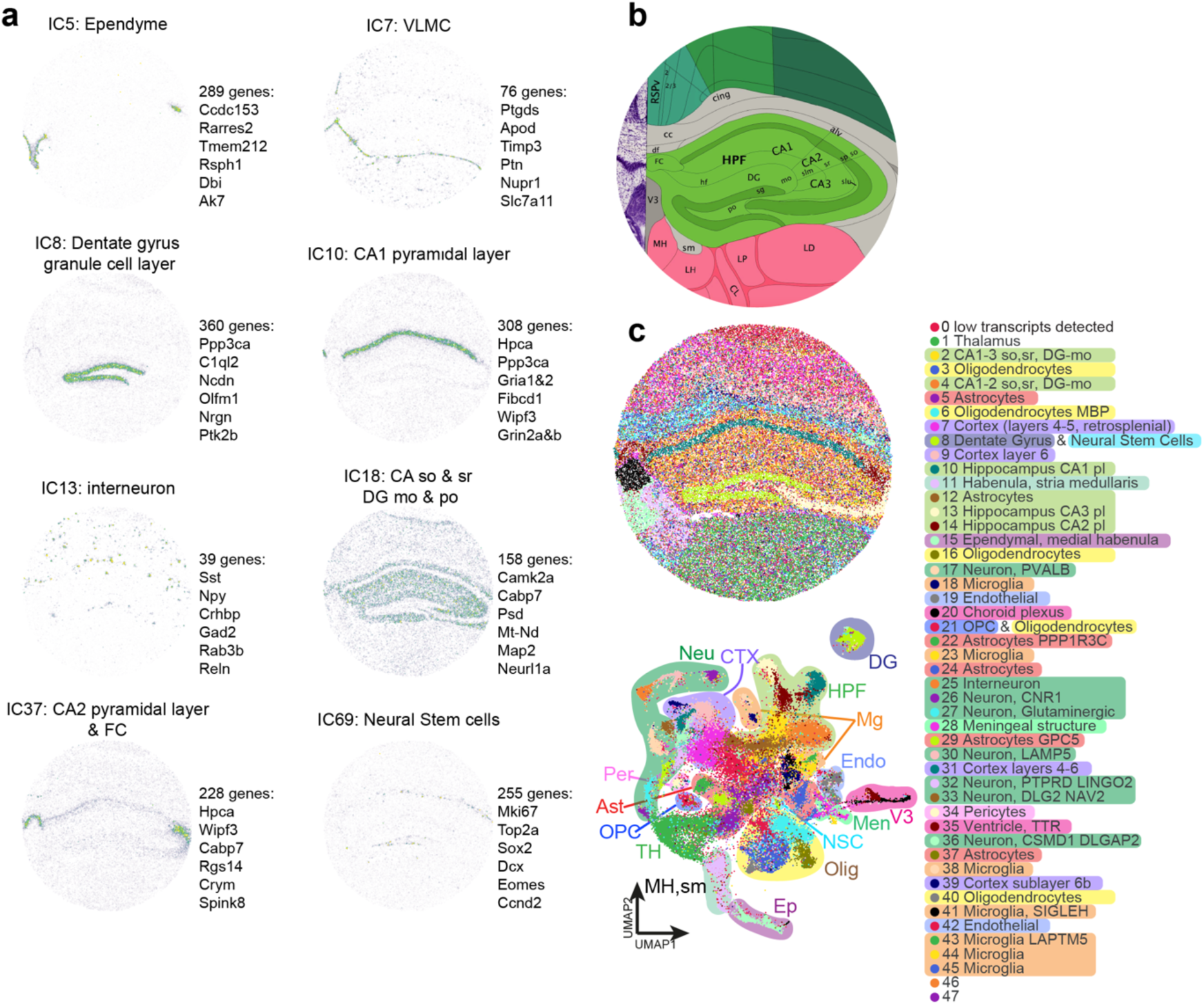
High resolution Slide-seqV2analysis using CFS allows for fine substructure definition and signal deconvolution. (a) Allen brain atlas reference for region of interest. (b) Spatial (top) and UMAP (bottom) projection of spots clustered using Louvain algorithm at a 0.95 resolution using all filtered and annotated ICs as input with detailed and broad (shading) cluster annotations. (c) Sample spatial projections of substructure- or cell type-associated ICs with their 6 most contributive genes (see Supp table 8 for full list of gene contribution by IC and Supp table 7 for IC annotations) Ast : Astrocytes, CA : Ammon’s horn, CTX : Cortex, DG: Dentate gyrus, Endo: Endothelial, Ep: Ependyme, FC: Fasciola cinerea, HPF: Hippocampal formation, Men: Meningeal substructure, Mg: Microglia, MH: Medial habenula, mo: Molecular layer, Neu: Neuron, NSC: Neural stem cell, Olig: Oligodendrocyte, OPC: Oligodendrocyte progenitor cell, Per: Pericyte, po: Polymorph layer, sm: Stria medullaris, so: Stria oriens, sr: Stria radiatum, TH: Thalamus, V3: 3rd Ventricle, VLMC: vascular and leptomeningeal cell.

### ISH-based ST data analysis

ISH-based ST approaches are emerging as powerful technologies for detailed characterization of transcript expression. Commercial solutions are now available for the detection of hundreds to thousands of gene transcripts via multiplexed panels at subcellular resolutions with centimeter-scale capture areas, leading to the generation of enormous datasets with considerable computational and analytical challenges. One such challenge is the identifications of regions of interest (ROI) with a more manageable size for detailed analysis. We thus propose the use of CFS for FISH-based ST screening and ROI identification by signal binning. To that end, we tested the approach on both Vizgen MERSCOPE Mouse Brain Receptor Map and Nanostring CosMX FFPE human non-small cell lung cancer reference datasets.

ISH-based ST analysis with CFS requires a user-defined pseudo-spot generation step to ensure compatibility with the standard CFS workflow. This step was conducted as described in the Methods section and illustrated in Figure 3a. The choice of the pseudo-spot area size aims at striking a balance between signal resolution and computational cost. As show in Figure 3b, IC kurtosis distribution, used as a descriptor of IC super Gaussianity, decreased with the increases in pseudo-spot size, and their number. We found that for the MERSCOPE mouse brain dataset 40 µm bin sizes (1600 µm^2^ bins) stroke the right balance between resolution and computational resources use for pre-screening application (Fig. 3b). Interestingly, the total number of IC-defining genes was observed to increase with decreasing pseudo-spot sizes, ranging from 219 to 302 unique contributing genes, with 10 and 450 µm bin sizes respectively, out of the 649 probed genes for this dataset (Fig. 3c). Without surprise, a higher resolution increased the number and resolution of clusters (Fig. s4). Most bin sizes between 20 and 90 µm appeared usable while, interestingly, a 10 µm resolution appeared unstable and over-clustered in addition to being computationally intensive.

**Figure 3.**
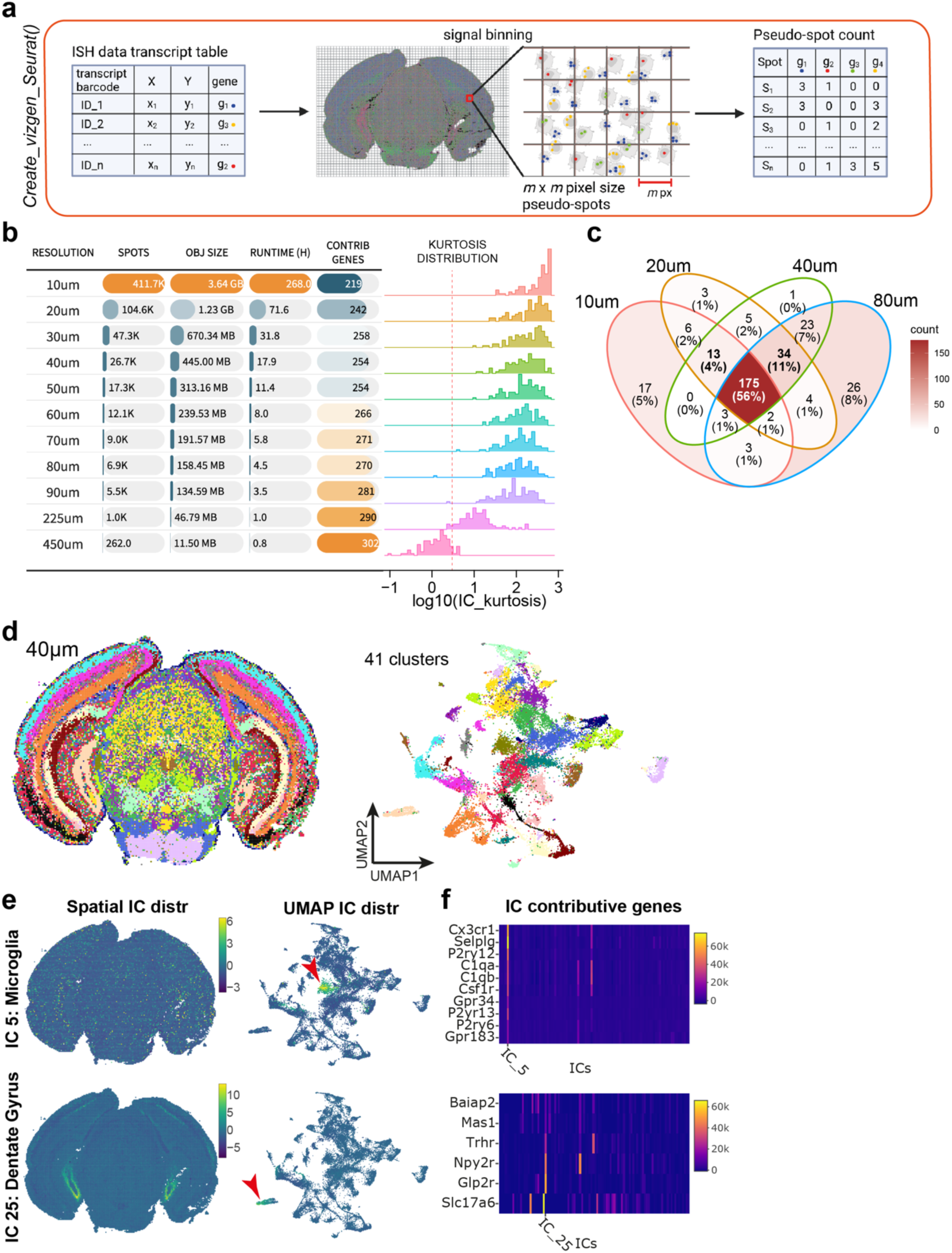
CFS allows for the analysis of ISH-based spatial transcriptomics data at varying degrees of resolution. (a) Schematic of the methodology behind the ‘*Create_vizgen_Seurat()’* function of CFS which generates pseudospot of user-defined m x m pixel size containing transcripts tabulated by the MERSCOPE technology to generate a count matrix input for the initiation of a Seurat object to input into the CFS Shiny application. (b) Recapitulative table of the impact of bin size m on object size, processing time, number of total contributive genes and kurtosis distribution of ICs (See Supp figure 4 for spatial and UMAP projections at different resolutions). (c) Venn diagram of contributive gene showing that 92% or more of contributive genes are detected in at least 2 levels of resolution with 225/254 (88.6%) contributive genes at m = 40 µm found in 3 or all 4 resolution levels (bold). (d-e) Sample analysis using CFS’s Shiny application for ICA signal deconvolution and annotation. (d) Spatial (left) and UMAP (right) projection of 40×40 µm pseudospots clustered using Louvain algorithm at a 1.0 resolution using all filtered and annotated ICs as input, generating 41 distinct pseudospot clusters. (e) Examples of spatial (left) and UMAP (center) projections of ICs associated to microglia (IC 5, top) and dentate gyrus substructe (IC 25, bottom) showing their distinct localization on the UMAP space, suggesting this level of resolution is sufficient to limit cell mixtures within pseudospots and capture specific cell populations. (f) Heatmap of contributive genes associated to the ICs in (e) from the 649 genes probed in the MERSCOPE experiment.

Despite the relatively low number of genes probed in FISH-based approach compared to whole transcriptome sequencing approaches, CFS was able to isolate structure and cell type-specific signal in the analyzed samples. For instance, the MERSCOPE mouse brain dataset at a 40 µm bin size yielded 100 leptokurtic ICs (min 12,78, max 588.60, Fig. 3d, Table S9-10), of which manual curation retained 63 with highly specific signals for cell types – IC 4: astrocytes, IC 5: microglia (Fig. 3e top), IC 14: endothelial cells – and brain ontologies – IC 8: pons, IC 25: dentate gyrus granule cell layer (Fig 3e bottom), IC 87: cerebral aqueduct (Table S9). The use of a limited set of targeted genes in ISH-based methods as opposed to whole transcriptome thus does not appear to impair the ability of ICA to identify cell- and ontology-specific signal (Table S10). The interpretation of the ICs can however be more imprecise based on the limited number of contributive genes (Fig. 3f).

The distribution of publicly available cancer datasets from the CosMX SMI platform by Nanostring also allowed for the evaluation of the modified CFS pipeline on FISH-based non-structured tumor tissue to assess the ability of ICA to deconvolute spatial signals of a more restricted dimensional nature (980 probed genes). The CosMX NSCLC dataset is comprised of multiple samples from distinct presentations of non-small cell lung cancer from different donors (5 donors, 8 samples) allowing for integrated sample analysis using the CFS workflow. Simply, samples were first binned as described previously (Fig 3a, Methods) using the *Create_CosMX_seurat* function with a 200×200 px (24×24 µm) pseudospot resolution. All 8 Seurat objects were then merged into a single 114 724 pseudospot object and processed simultaneously with the pre-processing pipeline (See Methods). All 100 ICs obtained passed the kurtosis threshold and 81 were kept following manual annotation (Table S11-12). Of the 980 genes assayed, 355 were used to define annotation-filtered ICs as contributive genes (36.2%). The filtered ICs describe common parenchymal and immune signatures for the tumor stroma, and common tumor programs between samples while uncovering phenotypical variations between tumors, as described by sample-enriched ICs (fig. s5).

Pseudospot clustering and annotation allowed the identification of multiple immune and stromal populations along with cancer-specific clusters (fig. 4a). Without surprise, while most cancer clusters were found to be patient-specific, immune and stromal clusters were found in more than one patient, such as IgD and IgG secreting B cells, macrophages, fibroblasts and erythrocytes (fig. 4b, s6). Of note, a neutrophil population with high expression of olfactomedin 4 (OLFM4) was found to be present in all samples. Spatial projection of these clusters (fig. 4c) allows for the rapid identification of these shared (or distinct) populations of interest for further analyses at a transcript level with other software solutions.

**Figure 4.**
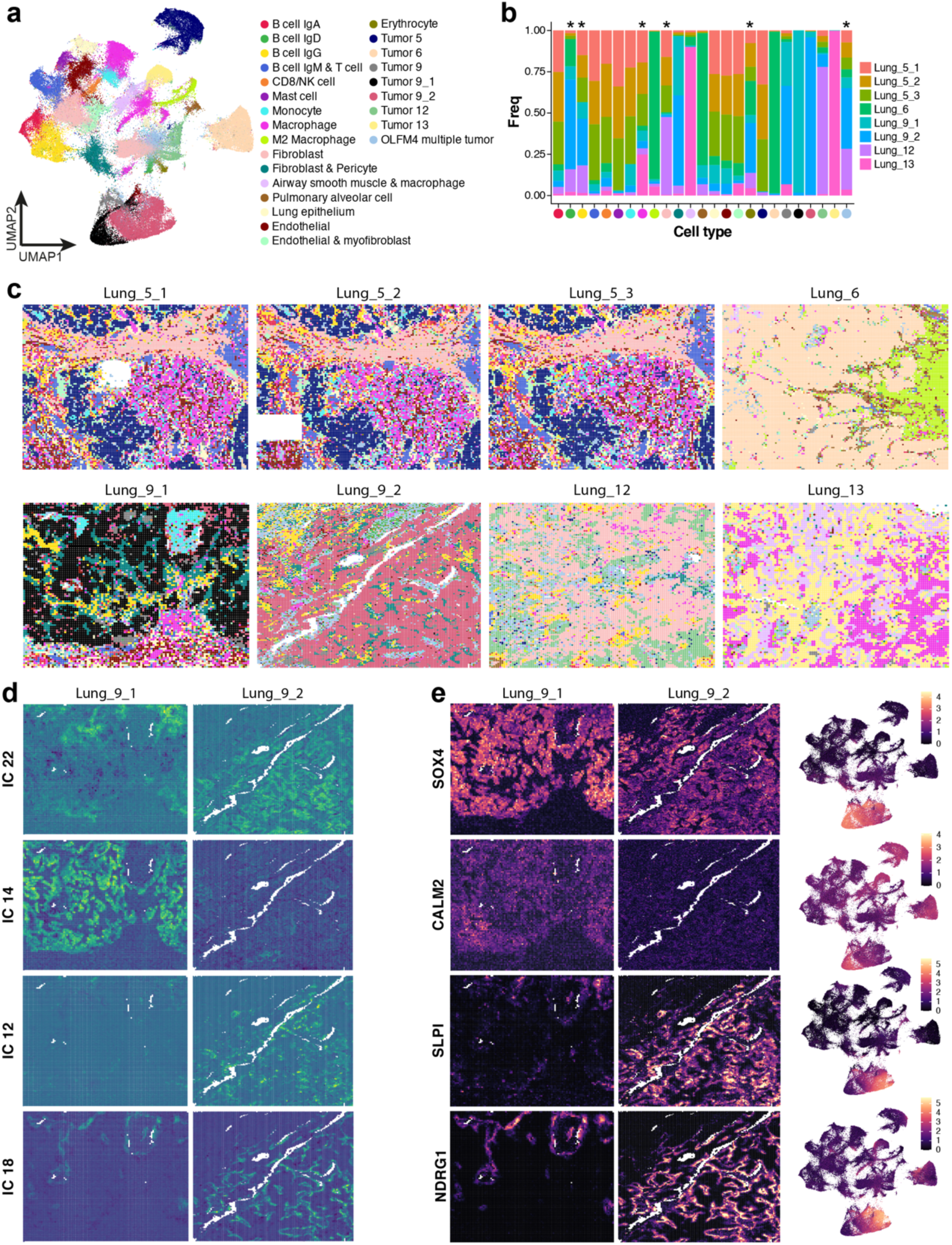
CFS enables the integration of multiple datasets to identify common and sample or patient specific gene signatures. After applying pseudospot generation methodology and sample merging of the Nanostring CosMX Lung cancer dataset, the combined object was analyzed using the standard CFS pipeline. (a) UMAP projection of 24 x 24 µm pseudospot clustered using Louvain algorithm at a 1.0 resolution and annotated using top contributive ICs (see Fig. S6 for UMAP coloring by sample). (b) Stacked bar plot illustrating the distribution of clusters between samples. * indicate clusters with >25% of pseudospots found in multiple patients (note that patient 5 has 3 samples and patient 9 has 2 samples). (c) Spatial distribution of pseudospot clusters calculated in (a) showing the relative composition in shared (i.e. B cells IgG : yellow, Macrophages : magenta, OLFM4 multiple tumor : light blue grey) vs sample-specific clusters, notably in both samples of patient 9 (Lung_9_1 & Lung_9_2, bottom left). (d-e) Detailed interrogation of common and distinct transcriptomic signatures between samples of patient 9. (d) Spatial projection of 4/9 Lung_9 cancer-associated ICs (see Supp table 11 for all IC annotations) on sample Lung_9_1 (left) and Lung_9_2 (right). (e) Spatial (Lung_9_1 : left, Lung_9_2 : center) and UMAP (right) projections of top common (SOX4, top) and sample-specific genes (CALM2, SLPI, NDRG1, bottom 3) as determined by IC contribution and differential gene expression analysis between clusters defined in (a).

Interestingly, CFS demonstrated ability to characterize intra and inter patient tumor heterogeneity when performing multi-sample integration analysis: While samples from different patients were well distinguished in the UMAP projection, cancer-associated pseudospots from both samples from patient 9, taken in different parts fo the same tumor, clustered closely while remaining locally distinguishable (fig. 4a, black and muted red clusters), allowing for the interrogation of the intratumoral heterogeneity in a spacially resolved manner. Both tumor sections show IgG B cell infiltration and fibroblast & pericyte stromal components (Fig. 4c yellow and dark teal clusters). Using ICA, we identified 9 ICs associated to the cancer component of these tumors with some shared between both samples (ex. IC 22) and others specific to one or the other (ex. ICs 12, 14 and 18, fig. 4d). Analysis of contributive genes to these ICs (Fig. 4e) revealed that SRY-box transcription factor 4 (*SOX4*) is expressed exclusively in patient 9 by all cancer cells, but sample 9_1 is characterized notably by the expression of calmodulin 2 (*CALM2*) while sample 9_2 cancer cells are enriched in secretory leukocyte peptidase inhibitor (*SLPI*) and N-myc downstream regulated 1 (*NDRG1*).

### CFS Reference-free deconvolution performance assessment

Reference-free unsupervised signal deconvolution is becoming an important aspect of ST analysis due to the limitations of single-cell reference-based approaches. Some of these limitations include the lack of relevant high quality single cell atlases for pathologic conditions or difficult-to-process tissues, or cell capture bias leading to missing cellular identities. The performance of supervised approaches is also deeply dependant on the quality of the reference’s annotation. At the time of writing, according to the benchmark by Li et al^9^, the best published tool for reference-free deconvolution being STdeconvolve which uses an LDA modelling approach for an optimized K number of topics for each dataset. We thus used it as a reference to evaluate CFS’s signal deconvolution performance. On the highly structured mouse brain sample from 10X Visium, we obtained 45 high quality ICs using the CFS pipeline while STdeconvolve found the K=38 condition to be optimal. All STdeconvolve topics were recapitulated by one or more ICs both by spatial distribution and contributive gene weight (figs 5a left and s7a left respectively). In most cases, such as in topics 13 or 9 (fig. 5a blue and green outlines respectively), CFS was able to further decompose the signal into 4 and 2 major components, often with a higher spatial definition (fig s7b). Similarly, for the heterogeneous 10X Visium breast cancer dataset, CFS yielded 46 ICs to STdeconvolve’s 30 topics (fig 5b left), with CFS components often appearing more spatially defined (fig 5b blue and green outlines & fig s7c).

**Figure 5.**
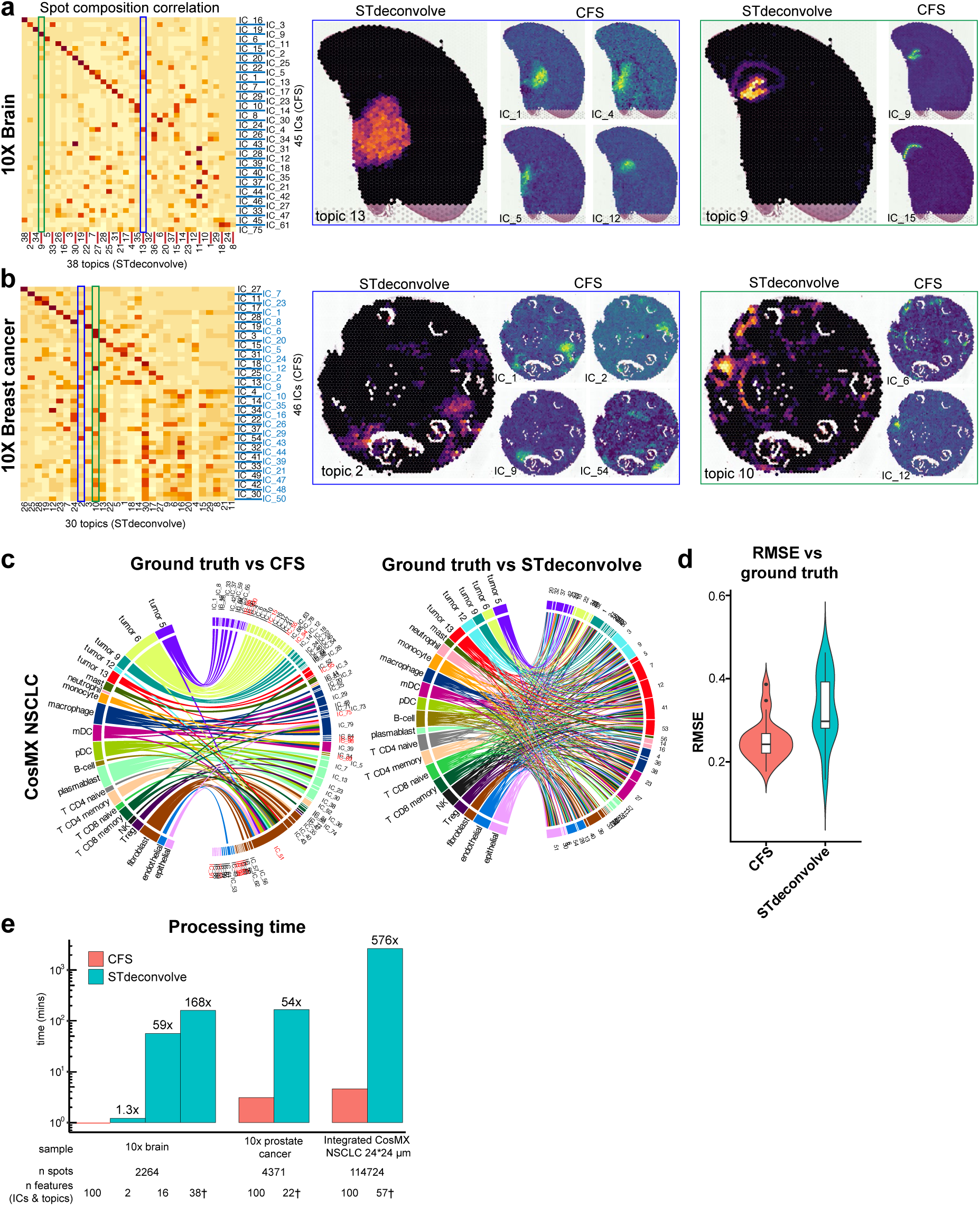
Comparative analysis of reference-free deconvolution results from CFS and STdeconvolve. (a,b) Correlation analysis based on feature-defining genes of reference-free deconvolution tools CFS and STdeconvolve (left) with 2 highlighted examples (blue and green outlines) of STdeconvolve topics vs CFS ICs for a specific structure in the 10X brain (a) and 10X breast cancer (b) samples. See supplementary figure S7 for more examples. (c) Gene signature signal correlation of CFS ICs and STdeconvolve topics vs. ground truth (left and right, respectively) displaying all Pearson’s correlations r values > 0.3. IC names in red indicate ICs that were rejected by manual annotation (fig 4), but were kept for unsupervised performance assessment, note IC_51 with high correlation with most cell types. See supplementary figure S8 for pseudospot compositional correlations (d) Root-mean-square-error (RMSE) of the deconvolved compositions using CFS or STdeconvolve (*n* = 112 822 pseudospots) compared to ground truth after ILR-transformation. One-tailed Diebold-Mariano *p* value 1.473 x 10^-5^. (e) Processing time comparison between CFS (red) and STdeconvolve (teal) shows the exponential benefit of CFS in processing time with increased dataset size. “n Features represents” the number of user-defined features (ICs for CFS, topics for STdeconvolve) for each processing run, with * indicating the optimal K value defined by STdeconvolve for this sample.

Comparative assessment of deconvolution performance was conducted using the CosMX NSCLC dataset with cell annotations as ground truth. Gene signature correlations showed that ICs generated with CFS tended to be associated to a unique cell type while all STdeconvolve topics correlated with multiple of cell types (fig 5c). Spot composition correlation of ICs and topics to the ground truth indicates that for both algorithms, most features correlate with a unique cell type (fig. s8). However, STdeconvolve topics showed a higher rate of mixed topics, i.e. topics associated with multiple cell types (9/57 vs. 4/81 for CFS ICs, fig. s8). With both methods however, most lymphoid populations (T CD8 naive and memory, NK and Treg) showed poor correlation with deconvolution features. Based on gene signature correlations, the root-mean-square error (RMSE) relative to ground truth of cellular proportions in each pseudospot, after ILR-transformation, was calculated for both CFS and STdeconvolve (fig. 5d). Results show that the CFS pipeline recapitulated the cellular composition with higher accuracy than using STdeconvolve topics (One-tailed Diebold-Mariano *p* value 1.473 x 10^-5^).

Finally, the computing efficiency and scalability of CFS’ ICA also proved to be a major improvement over STdeconvolve’s LDA method. Processing times for STdeconvolve for a single optimal K value ranged from 54 to 576 times longer than with CFS. For instance, processing the integrated CosMX NSCLC dataset (fig. 4) of 114 724 pseudospots with 980 probed genes took STdeconvolve 2657 minutes (1.85 days) to process vs. 4.6 minutes with CFS (benchmark does not consider the K optimization step of STdeconvolve).

## Discussion

In this paper we presented CellsFromSpace or CFS, a user-friendly and reference-free analytical framework for spatial transcriptomics data that leverages independent component analysis to deconvolute and integrate ST data. We have demonstrated that CFS extracts spatially distributed signatures as well as diffuse cell signatures.

The cornerstone of the CFS pipeline is the facilitated manual curation and annotation of ICs generated by the pipeline. While time consuming, this critical step not only allows for the elimination of non-specific noise, but also consists of the main data interpretation step. To facilitate and enhance this crucial step, we created an easy-to-use Shiny UI to provide any user (clinician, biologist, bioinformatician) with all the analytical and visualization tools to easily and confidently annotate and interpret the data generated with any major ST technology currently commercially available. Manual IC annotation enables the rapid and reference-free identification of cell types and states within samples and, among other things, the identification of genes with similar spatial distributions, defined as IC contributive genes. Manual curation also allows for the elimination of non-specific, over-specific, redundant, artifactual or uninterpretable ICs, thus denoising the dataset with a minimal loss of relevant signals thanks to the independent property of ICA. While we believe in remaining in the latent ICA space for data interpretation and analysis, CFS also implements the traditional workflow used in single cell transcriptomics and ST, which collapses the latent dimensions into clusters and two-dimensional projections for easy data visualization (UMAP, t-SNE, etc.) and downstream differential gene expression analysis.

ICA has seen a steady rise in popularity in the last few decades in a wide variety of application fields, each time with specific interpretations of the algorithm tailored to the signals and sources present. In the special case of ST data however, where ICs signal sources are defined biological entities (i.e. individual cells) our application seemed to reveal particular properties of ICA in the context of cellular biology, namely 1) the importance of signal sign, where the long tail of the IC distribution is associated to the cellular source identity, 2) relative positive ICs weights can directly be used as proxy for cellular abundance within sources (spots) for direct proportional deconvolution of signal. The latter enhancing ICA’s performance as a reference-free signal deconvolution approach.

Unsupervised reference-free signal deconvolution methods present considerable advantages over reference-based methodologies relying on single cell atlases. Beyond the independence from the availability of high quality reference datasets, reference-free methods are unaffected by single cell methods’ limitations such as compositional or transcriptional processing artifacts^29,30^ or the quality of single cell atlas annotations which can propagate cell misclassifications. Reference-free approaches are also useful for conditions for which single cell atlases might not be able to recapitulate unique phenotypes, such as for cancer cells with a high inter-patient heterogeneity. Finally, by avoiding the identification and processing of a high-quality reference dataset sample processing is made simpler and more accessible.

Performance assessment shows that CFS outperforms STdeconvolve with a higher specificity and log-scale acceleration of the processing time. Unlike STdeconvolve, CFS does not attempt to directly infer cell types proportions in each spot. We aim at minimizing assumptions regarding the extracted components as they do not always represent cell types but sometimes describe cell activities or cell-to-cell interactions and as such can represent complementary signal to cellular composition. This approach allows users, who are experts in their respective fields, the freedom to concentrate their ST data analysis on the signal they consider relevant.

Further development of CFS is currently underway for downstream analysis of IC interplay within tissues. Cross-correlation of 2D signal remains an unresolved area of data analysis with great potential for ST experiments in order, for instance, to identify co-localizing, mutually exclusive, or interfacing signals. For example, such analyses would be of great value to map cellular interactions between tumor and effector cells to identify potential mechanisms of tumor rejection or evasion and identify key molecular drivers of these interactions via ligand-receptor analyses.

## Conclusion

In this work, we presented a new framework and tool for the analysis of spatial transcriptomics data. CellsFromSpace is versatile with its support for all commercially available ST technologies, independent of high-quality reference datasets, easy to use with its Shiny UI, compatible with other single cell and ST analysis packages, and easily allows for the integrated analysis of multiple samples. We hope CFS will increase the accessibility and ease of ST data analysis to researchers and improve data interpretation.

## Software availability

CFS is an R package which can be downloaded in R using the devtools package from github directly at https://github.com/gustaveroussy/CFS. Tutorial, examples and documentation can be found at https://codimd.univ-rouen.fr/s/w0oZMV6fz.

## Acknowledgements

We acknowledge “une main vers l’espoir”, Imagine4Margo, and the Dell foundation that support our team through recurrent funding or donation. We acknowledge the bioinformatic and the genomic facilities of Gustave Roussy for the constant support to implement new technologies.

## Supplementary Figure legends

**Figure S1.**
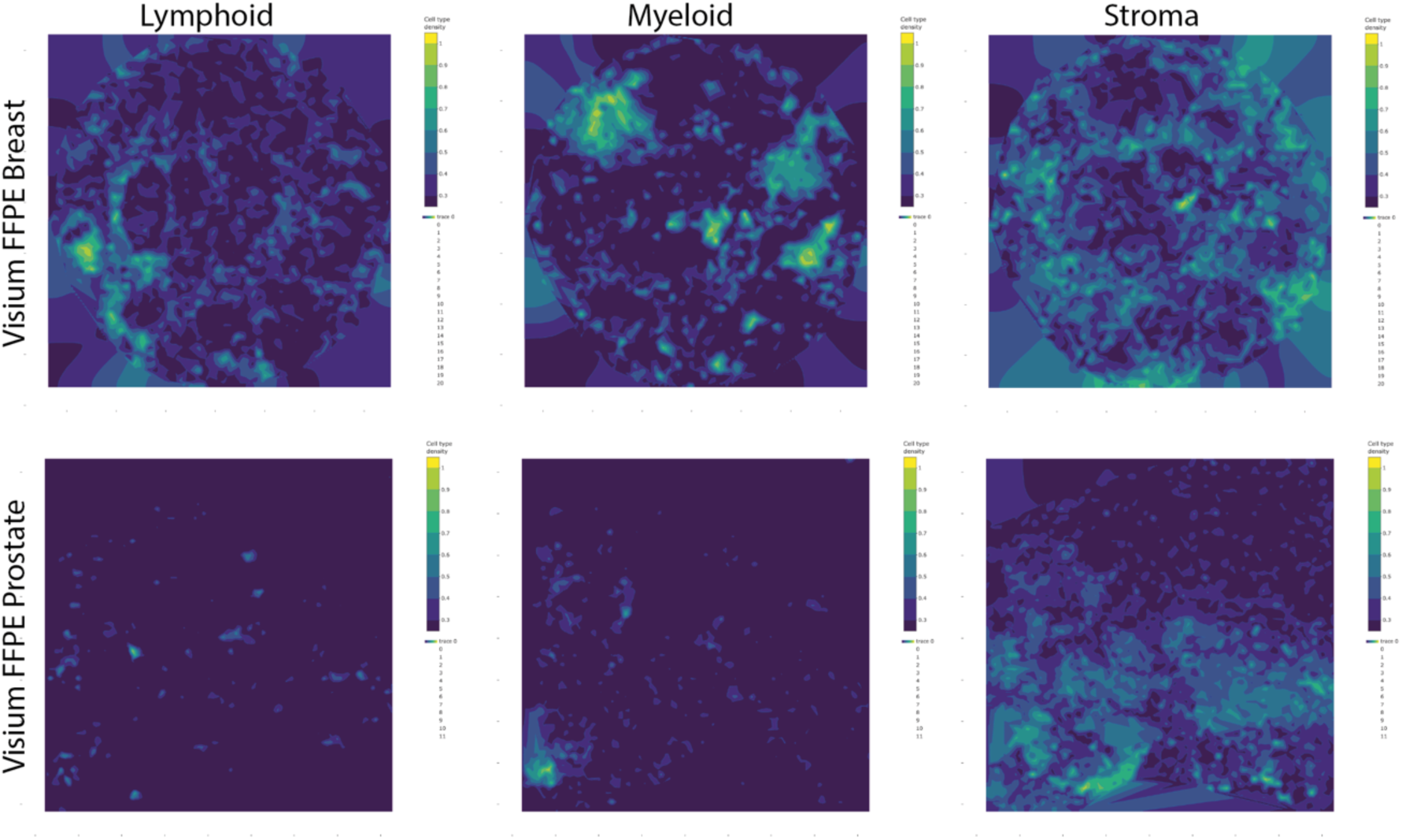
Signal density of tumor samples’ stromal components. Kernel density representation of the sum of lymphoid (left), myeloid (middle), and non-immune stroma (right) ICs in the Visium FFPE Breast cancer sample (top) and FFPE Prostate cancer sample (bottom) showing the global spatial localisation of each cellular component.

**Figure S2.**
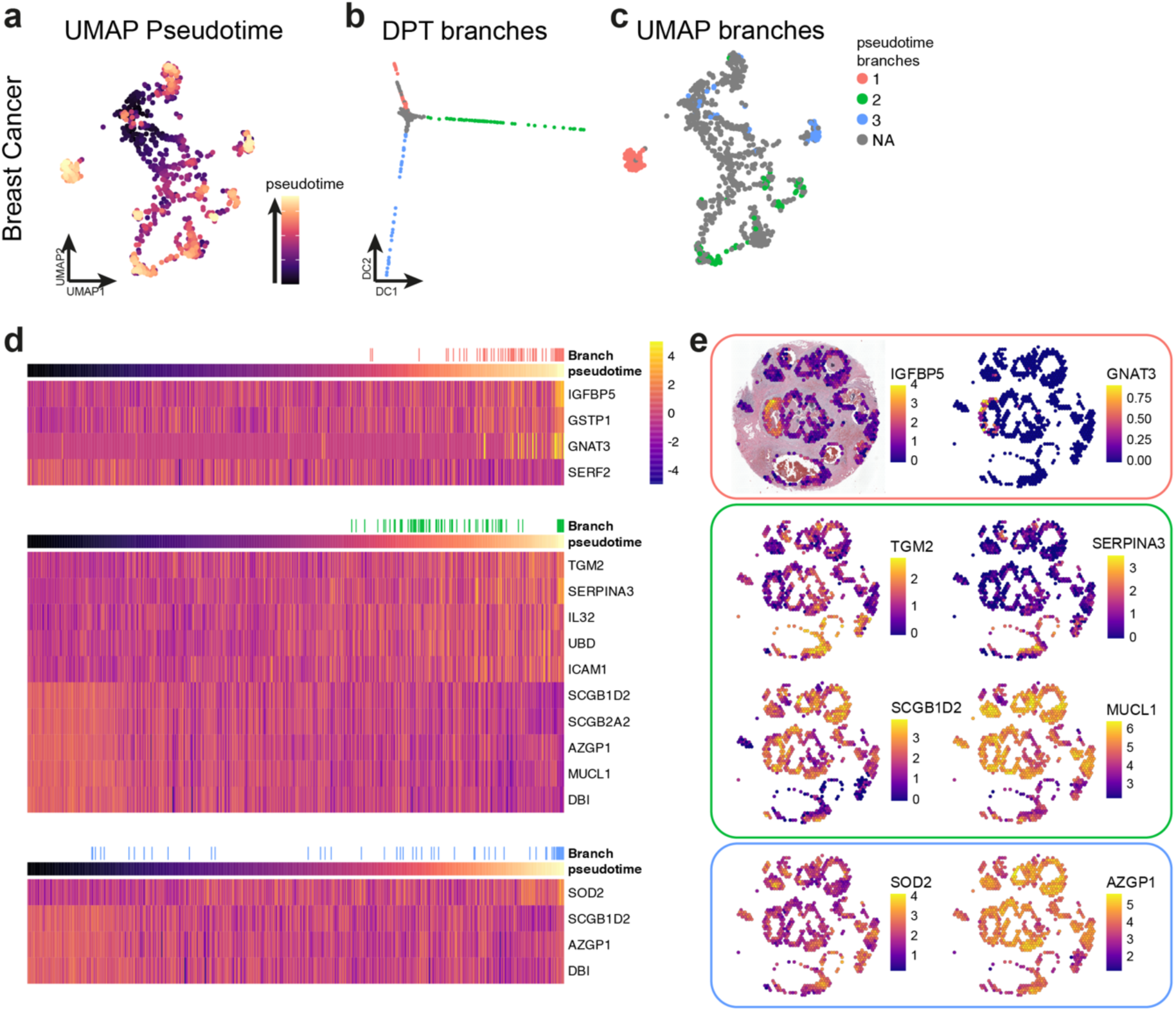
Trajectory inference of breast tumor’s cancerous regions. (a) UMAP projection of pseudotime calculated with Destiny’s DPT method. (b-c) projection of the distinct phenotypic branches identified by DPT on the first two diffusion components of the diffusion map calculated by DPT (b) and on the UMAP embedding calculated with cancer-related ICs (c). (d) heatmap representation of the expression levels of genes identified by glmnet for their association with branches 1 (top), 2 (middle) and 3 (bottom). (e) Spatial projection of expression levels of some of the genes identified in (d) for each branch.

**Figure S3.**
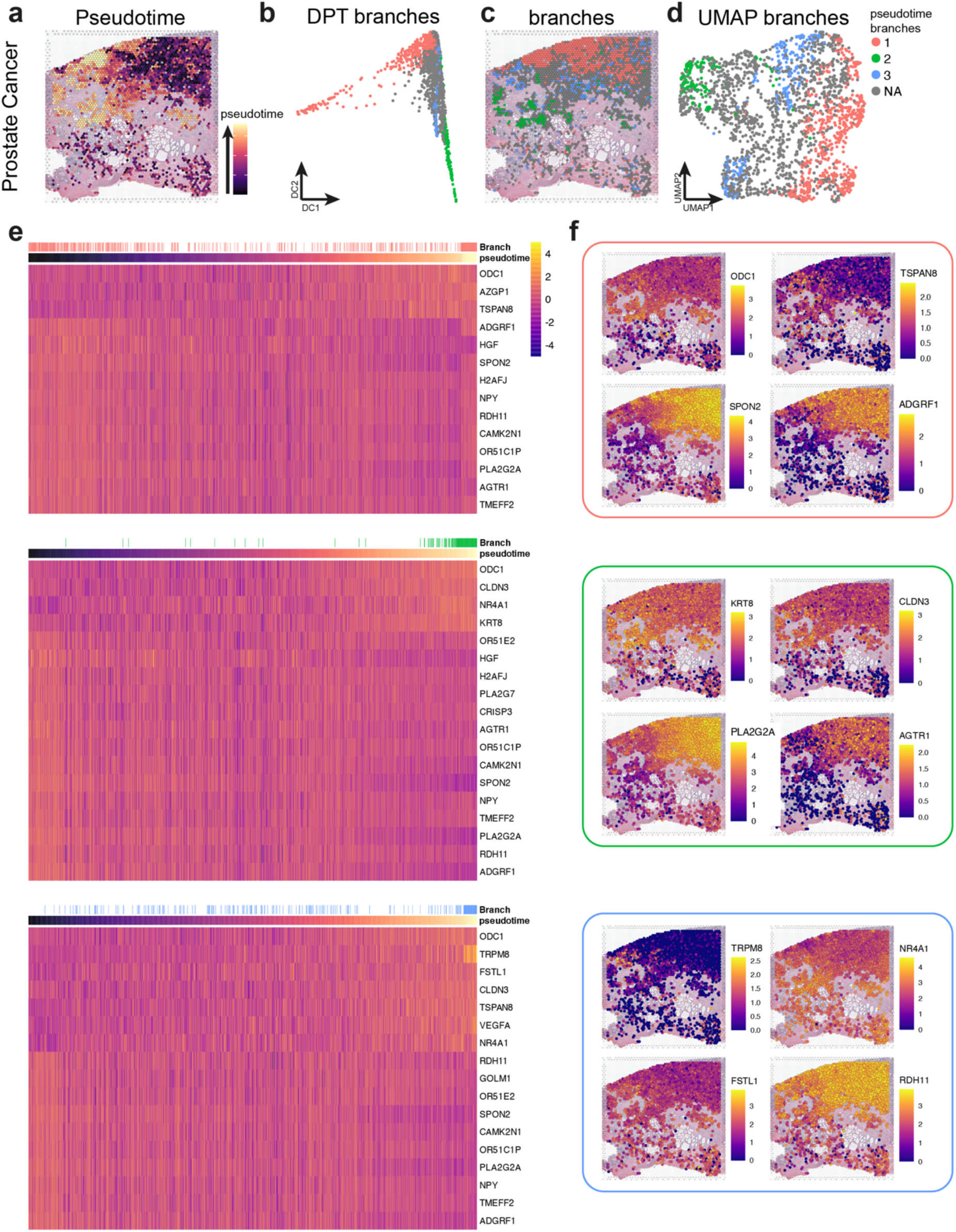
Trajectory inference of prostate tumor’s cancerous regions. (a) Spatial projection of pseudotime calculated with Destiny’s DPT method. (b-d) projection of the distinct phenotypic branches identified by DPT on the first two diffusion components of the diffusion map calculated by DPT (b), on the spatial distribution (c), and on the UMAP embedding calculated with cancer-related ICs (d). (e) heatmap representation of the expression levels of genes identified by glmnet for their association with branches 1 (top), 2 (middle) and 3 (bottom). (f) Spatial projection of expression levels of some of the genes identified in (e) for each branch.

**Figure S4.**
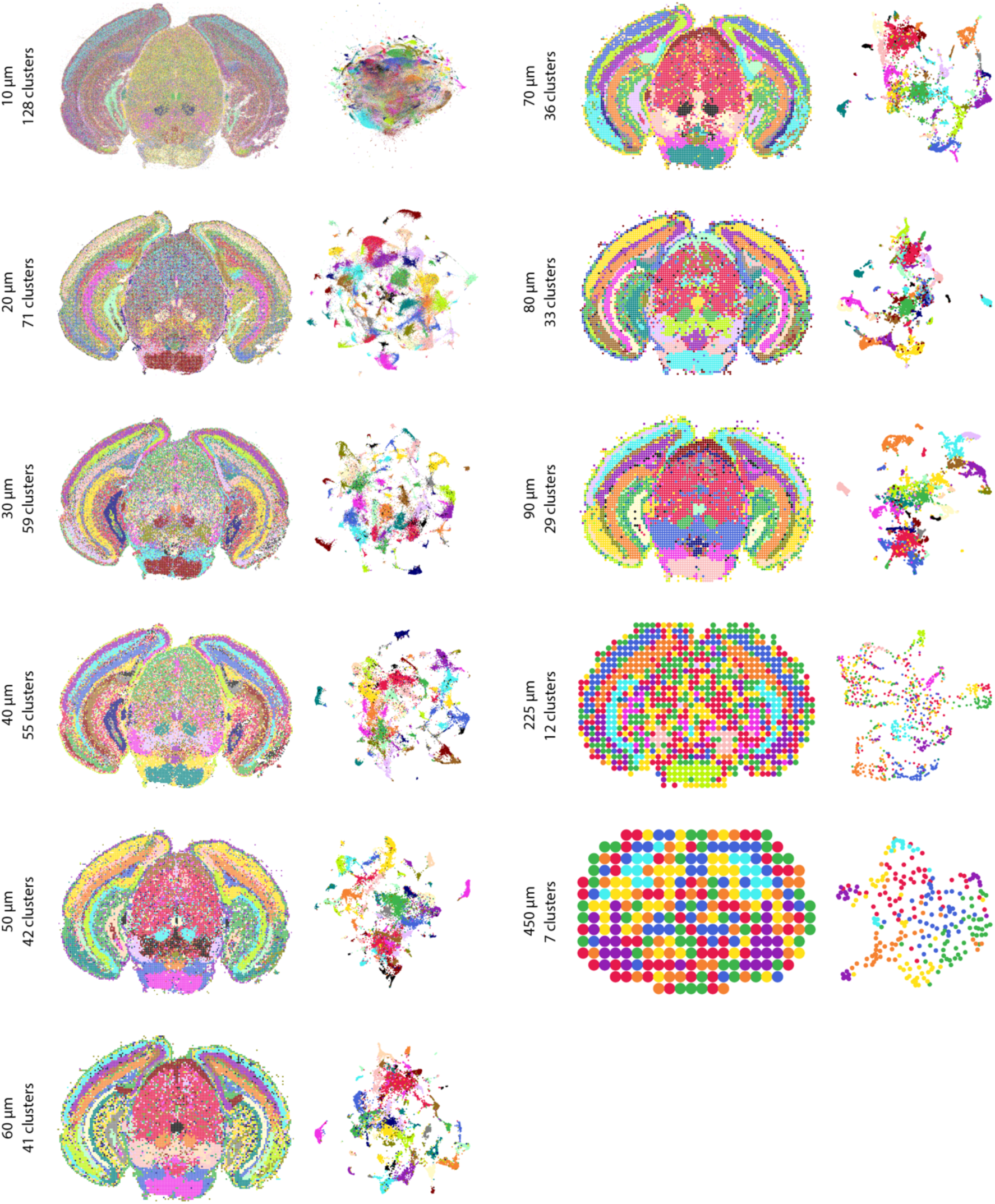
Impact of pseudospot size on resolution on clustering in ISH-based methods. Spatial (left) and UMAP (right) projections of pseudospot clustering at each pseudospot size evaluated. Clusters and UMAP embedding were calculated using all leptokurtic ICs for each condition with a constant louvain resolution of 1.2 for cluster identification and default UMAP parameters with a spread of 3 in all cases. The number of total clusters calculated is indicated for each condition. Note that the UMAP and number of clusters for the 40 µm bin size is different than presented in figure 3d because here all 100 ICs were used for calculation, while the calculations were done using 63 of those ICs after manual annotation and curation for the data presented in figure 3d.

**Figure S5.**
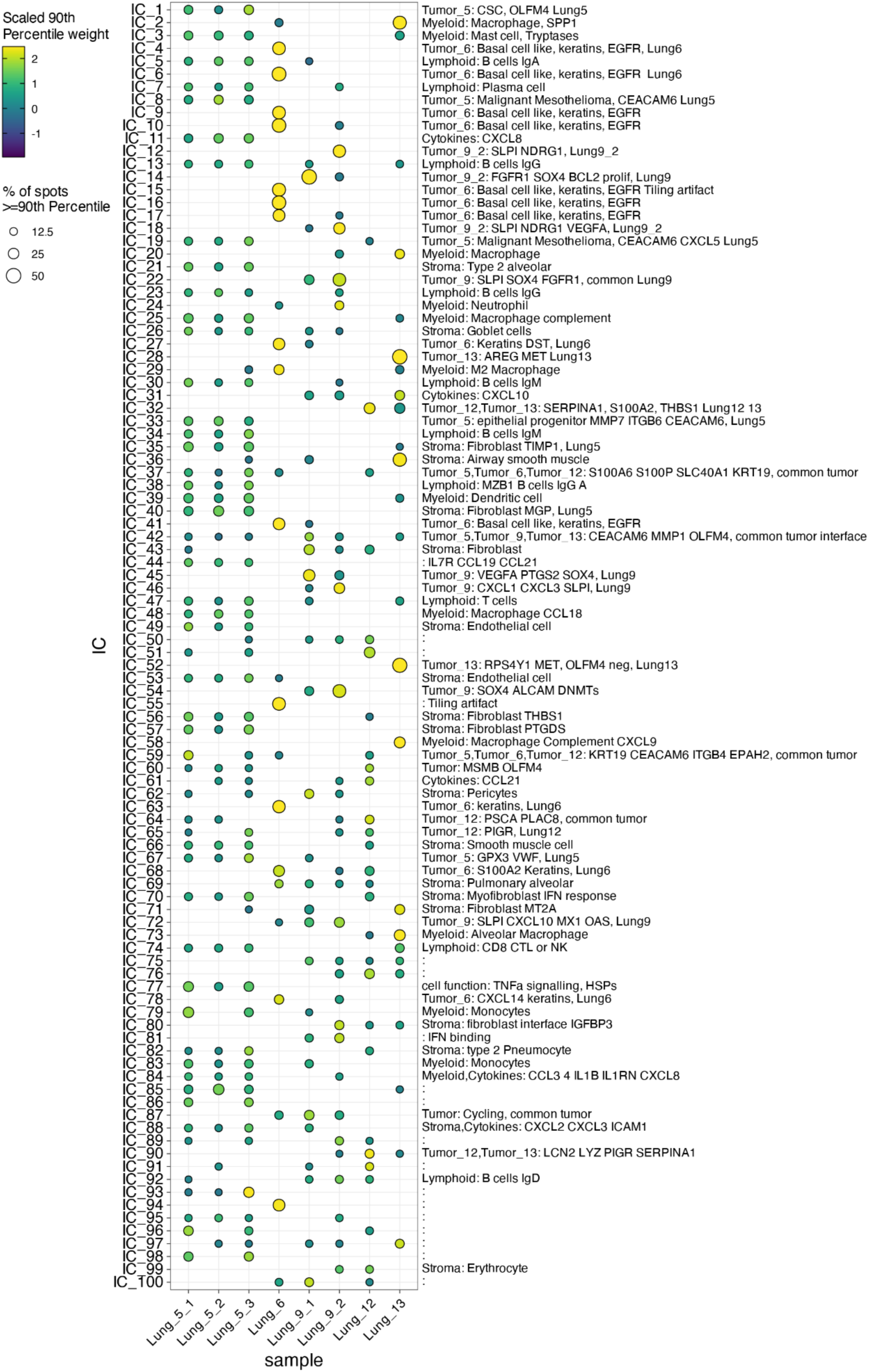
Sample distribution of ICs in the integrated CosMX Lung tumors dataset. Dotplot displaying the distribution of spots above 90th percentile weight value for each IC and the respective IC annotation (right). Color scale indicates the row-scaled value of the 90th percentile of IC weight for each IC per sample and point size is calculated based on the % of spots in each sample to be above the 90th percentile for each IC, with a minimum value of 10% below which points are censored for increased readability. ICs which are found in multiple samples (ex ICs 5, 13, 20, 26, 47, etc.) are often stroma-associated while sample- or donor-specific ICs are often cancer-related (ex. ICs 1, 4, 12, 28, etc.), but some cancer ICs were also found to be shared between samples (ex. ICs 32, 37, 42, 59, etc.). Note that samples 5_1, 5_2, and 5_3 are highly similar as they are serial cuts of the same tumor section, which explains why no sample-specific signal is found for this donor. ICs with no or incomplete annotations were filtered out in downstream analysis because they were either uninterpretable, redundant or artifactual.

**Figure S6.**
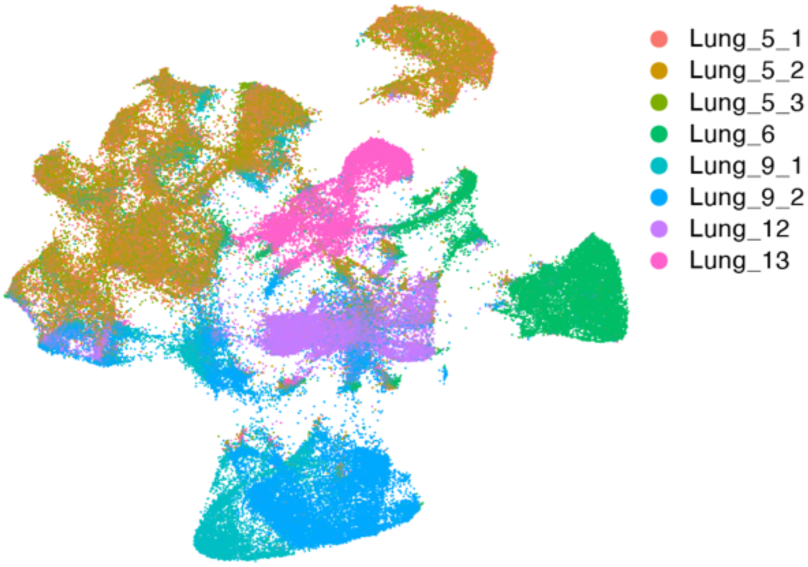
Sample of origin for each pseudospot of integrated analysis of CosMX Lung tumors dataset. UMAP embedding of all integrated pseudospots colored by sample of origin.

**Figure S7.**
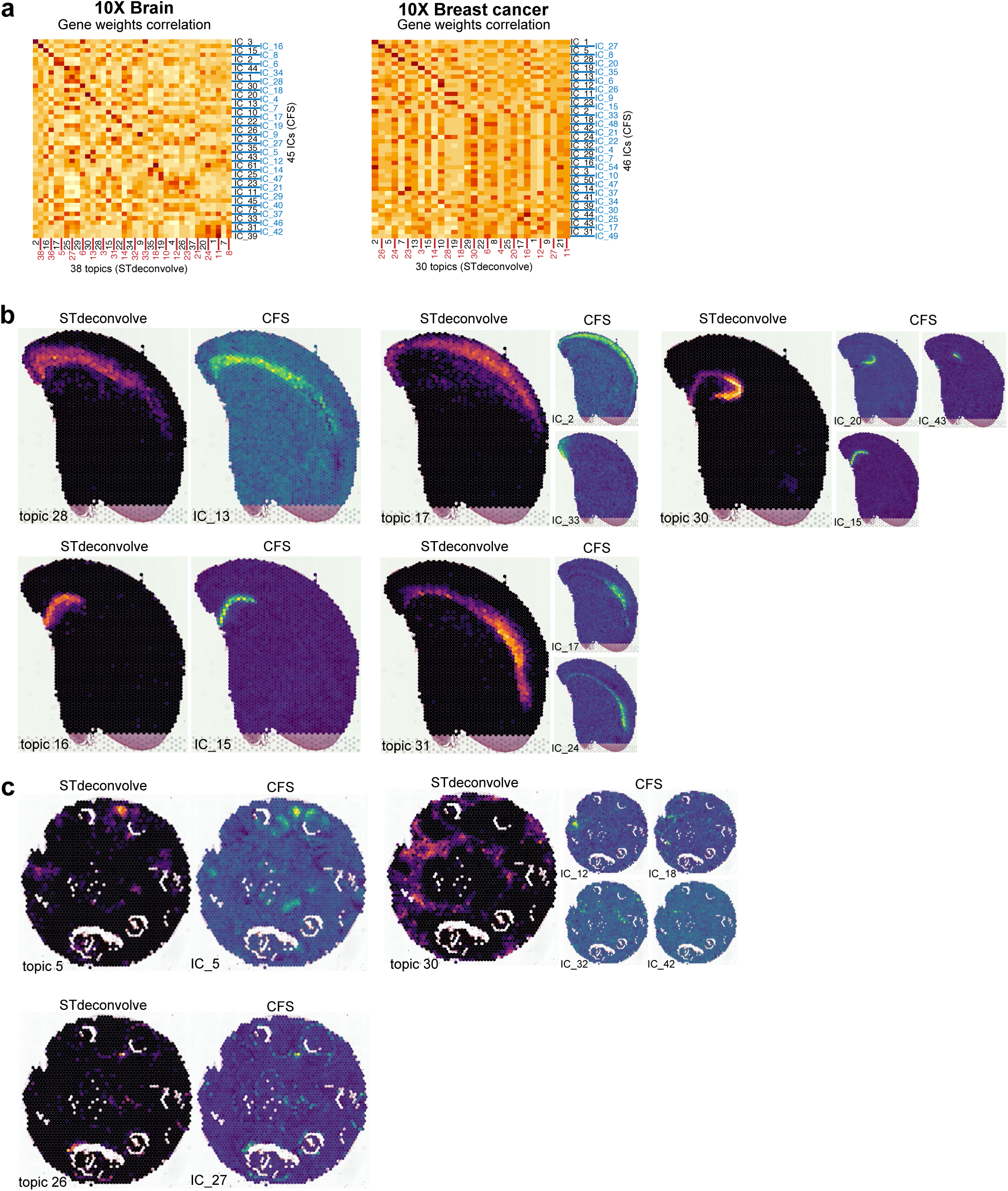
(a) Correlation of defining gene weight correlations between STdeconvolve topics and CFS ICs for the 10X brain (left) and 10X breast cancer (right) samples. (b-c) supplementary detailed examples of CFS ICs (blue-yellow scale) corresponding to interesting STdeconvolve topics (black-yellow scale) in 10X Visium samples of mouse brain (b) and breast cancer (c). Highlights the general ability of CFS to dissect topics into more ICs and for structure-related ICs to be less diffuse than associated topics.

**Figure S8.**
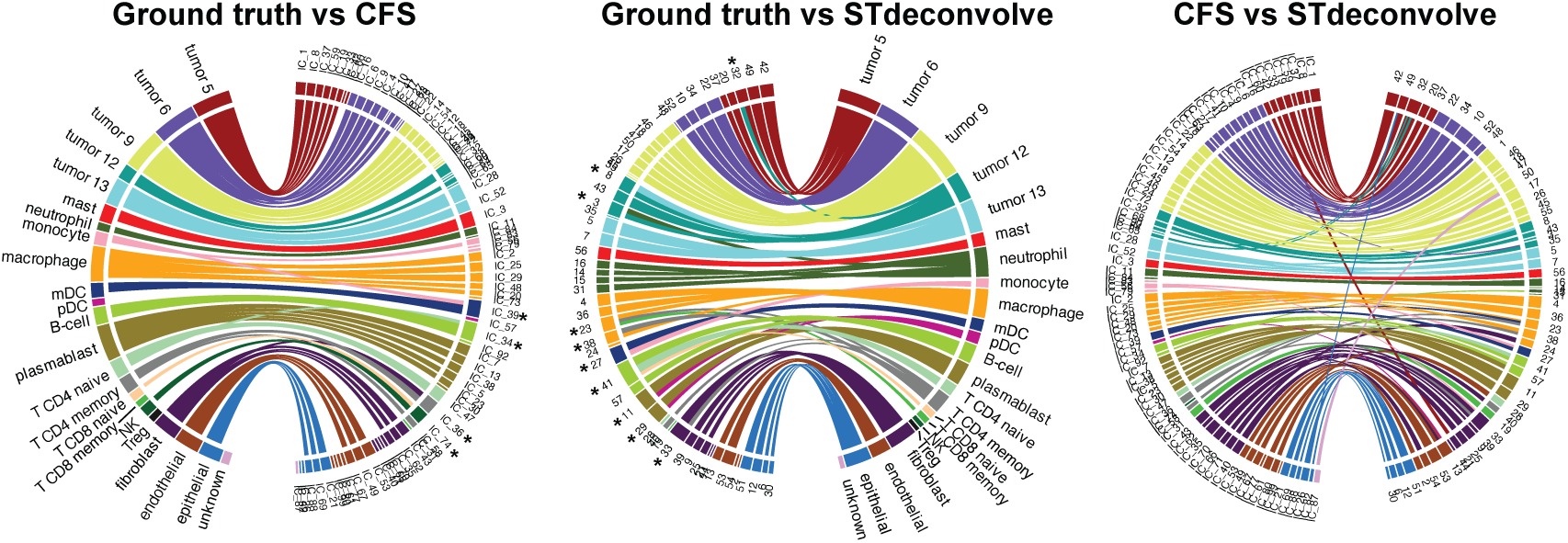
Pseudospot compositional correlation of CFS ICs and STdeconvolve topics vs. ground truth (left and middle, respectively), and CFS ICs vs. STdeconvolve topics (right) showing all Pearson’s correlations r values > 0.2.

## Supplementary Table legends

Table S1: Annotations of the independent components for the mouse brain Visium dataset.

Table S2: Most contributing genes to the independent components for the mouse brain Visium dataset.

Table S3: Annotations of the independent components for the breast DCIS Visium dataset.

Table S4: Most contributing genes to the independent components for the breast DCIS Visium dataset.

Table S5: Annotations of the independent components for the prostate cancer Visium dataset.

Table S6: Most contributing genes to the independent components for the prostate cancer Visium dataset.

Table S7: Annotations of the independent components for the ssHippocampus Slide-seq V2 dataset.

Table S8: Most contributing genes to the independent components for the the ssHippocampus Slide-seq V2 dataset.

Table S9: Annotations of the independent components for the mouse brain MERSCOPE dataset.

Table S10: Most contributing genes to the independent components for the mouse brain MERSCOPE dataset.

Table S11: Annotations of the independent components for the integration of the lung cancer COSMX dataset.

Table S12: Most contributing genes to the independent for the integration of the lung cancer COSMX dataset.

